# An algorithm for drug discovery based on deep learning with an example of developing a drug for the treatment of lung cancer

**DOI:** 10.1101/2023.05.07.538843

**Authors:** Dmitrii K. Chebanov, Vsevolod A. Misyurin, Irina Zh. Shubina

**Affiliations:** Department of Molecular Biology of Cancer, BioAlg Corp; The Russian Melanoma Professional Association (Melanoma.PRO)

## Abstract

We have designed an algorithm implemented in a software platform for the development of new anti-tumor drugs in the form of small molecules. Several molecules were generated for the treatment of patients with lung cancer as an example. At the initial stage, we identified the targets for the therapy. Firstly, we evaluated the expression profile of the genes most associated with poor clinical outcome in patients with lung cancer using deep learning. Additional patients’ data were gained by generative adversarial neural networks (GAN) technology. As a result, a set of genes was successfully selected, which expression was associated with poor prognosis. We identified the genes that could distinguish normal tissue from tumor tissue using another deep learning model that was trained to predict normal and tumor tissue based on gene expression. The other genes were considered as targets for targeted lung cancer therapy. After that, a module was developed that predicts the interactions of inhibitors with proteins. For this purpose, the amino acid sequences of proteins were represented in vector form, as well as formulas of chemical compounds interacting with proteins. In addition, a deep learning-based module was developed that predicts the IC50 in experiments on cell lines. Virtual pre-clinical trials with the selected inhibitors were performed to identify relevant cell lines for laboratory experiments. As a result, the study obtained formulas of several molecules with the predicted binding to certain proteins.

## Introduction

The search for new agents for treatment of patients with cancer is still a vitally important issue due unresolved problems of relapses and drug resistance to antitumor therapy. The challenge is that it is necessary to develop new drugs which should be more effective than those already registered. It leads to the enhancement of the cost for the research and production of the new drugs.

The computer approach allows us to solve this problem of creating new drugs. Deep learning technologies are successfully used in many areas of science and industry giving an opportunity to solve the problem with an unprecedented level of abstracting, inaccessible to human mind.

The source of new drugs may include a machine learning model trained to predict the therapeutic properties of molecules and used to search in the library of chemical compounds. Machine learning can be also used to predict drug-protein interactions, which will make it possible to identify well-targeted proteins, as well as molecules that are candidates for inhibitors. Another possibility of the machine learning model is the prediction of the result of the IC50 experiment taking into account the genomic expression profile of the cell line and the molecule structure. Such a model can predict whether the experiment with the cell lines and molecules can reach the established IC50 value, thereby emulating the cellular experiment in the *in silico* mode.

Our study has demonstrated that machine learning helps to solve the key problems of drug discovery, such as target identification in a tumor, followed by the selection of potential molecular inhibitors for the targets, as well as the design of preclinical studies and the experimental proof of the concept.

### 1. Target identification

#### Introduction

The discovery of new drugs is a process that mostly depends on the correct identification of targets. Approaches for target identification should take into account the following: 1) targets should be specific and determine the differences between the tumor and normal tissue; 2) targets should mediate tumor cell survival; 3) be available for small molecule therapy.

We believe that the most promising targets are the mutant genes (genetic disorders) that are responsible for a more aggressive disease associated with the reduced overall survival and relapse-free interval. Thus, the task of target identification is confined by the problem of machine learning for the selection (identification) of genes that are essential for successful prediction of overall survival and relapse-free interval.

At the same time, these genes should have different expression levels compared to that of normal tissue. Regarding these conditions the developed new drugs will have a significant effect on tumor cells and be relatively less toxic to normal cells.

#### Materials and Methods

To assess the effect of gene expression on disease prognosis, we used the data of gene expression from the open database TCGA https://gdc.cancer.gov (1). Gene expression (RNA-Seq) data were downloaded, then the values were normalized to the expression levels of the control or so-called “housekeeping” gene GAPDH for further updating the database with the new data. The data of overall survival (OS), progression free interval (PFI) and the same parameters within the follow-up period were derived from this database.

We studied lung cancer as an example diagnosis in 514 patients.

To reduce noise during training, we selected only those genes that are included in tumor-associated signaling pathways according to the KEGG resource in total 1821 gene (2). The problem of OS prediction was successfully solved in our previous study (3). Similarly, the target variable for prediction was the possibility of the patient to surpass the median PFI or OS for the whole dataset. We used the Keras library for deep learning.

Firstly, we trained the algorithm on the available data of 514 patients, but mean ROC-AUC after 5-fold cross validation was 0.69 (0.61-0.74) for OS and 0.61 (0.54-0.69) for PFI, which indicates the unsatisfactory quality of the prediction. The reason is that that was a too small dataset for the application of neural networks, even taking into account more than 1800 features. To avoid this, we obtained additional patients’ data using generative adversarial networks (GAN) technology, which has been successfully used in various industries, such as image generation (4). To implement GAN, we used the sdv library (5), in particular, the CTGAN module (6).

#### Results

We generated 50k patients’ data to reach the ROC-AUC value equal to 0.73. We used the Lasso linear regression algorithm with 5-fold cross validation to determine the significance of the features. The result of each experiment was obtained as a list of genes ordered by decreasing impact on the effect. We intersected the lists of genes obtained in the experiments with OS and with PFI.

As a result, 36 genes were found which expression was associated with impaired survival parameters of patients with lung cancer. Table 1 presents some characteristics of these genes.

**Table 1.**
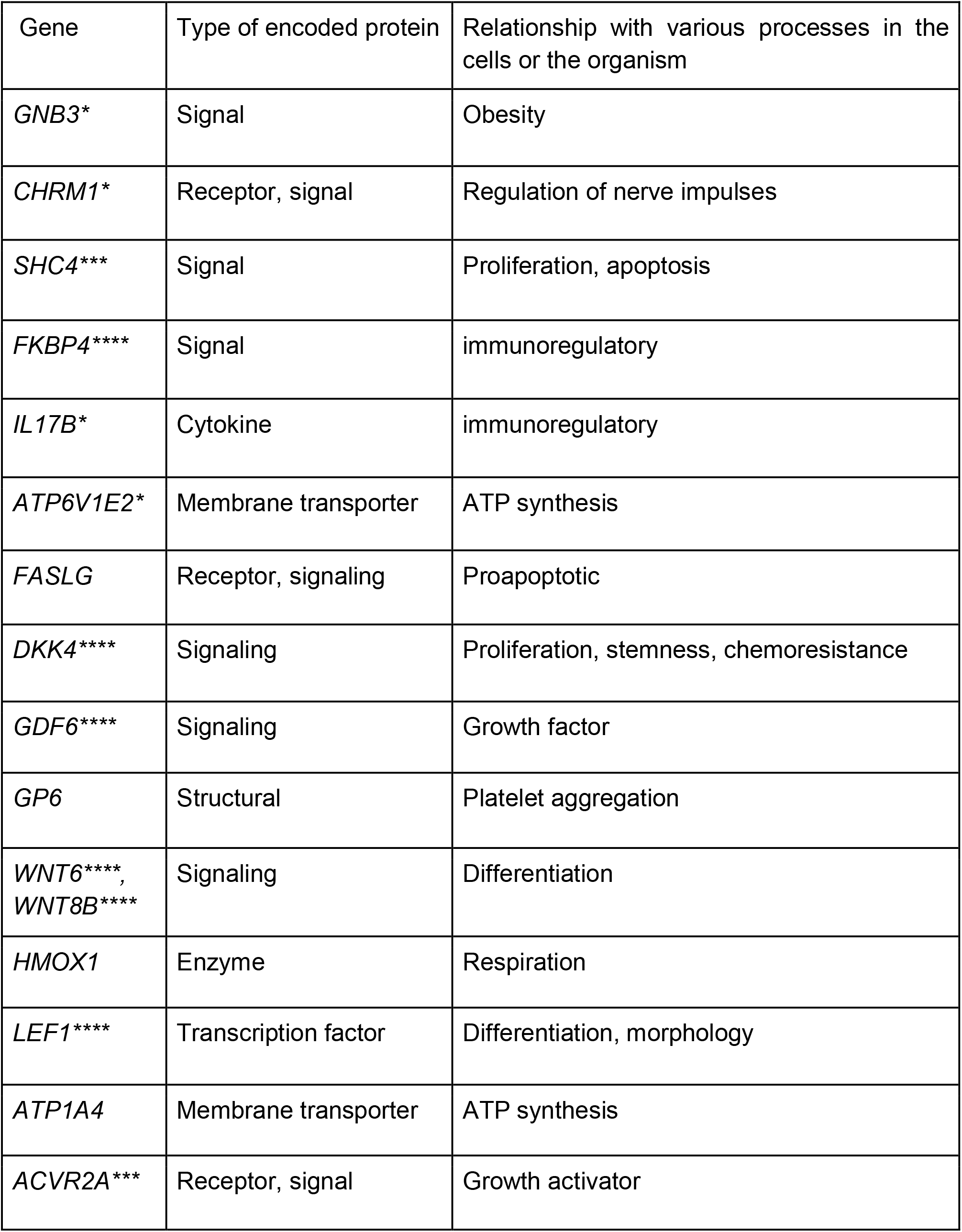
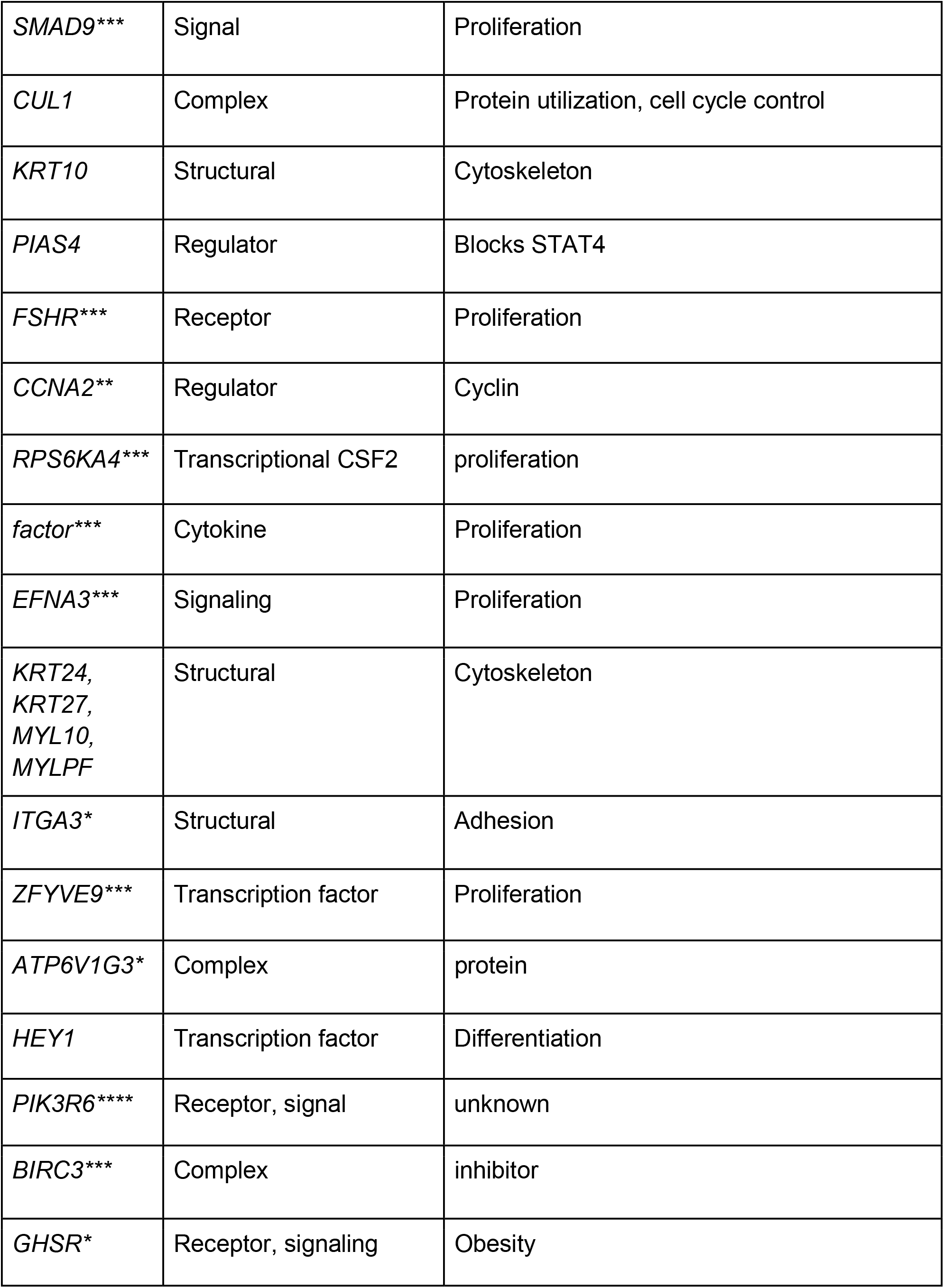
Biological features of the discovered genes.

### 2. Tumor-normal filter

#### Introduction

A number of the identified genes are responsible for the processes of energy and synthesis of proteins that are equally important for both tumor cells and normal cells. Accordingly, their inhibition will negatively affect not only tumor cells, but also the organism as a whole. Therefore, we used one more approach for the selection of target genes. We used the data of tumor and normal tissue gene expression to exclude the genes which expression is comparatively different in normal and tumor tissues. An additional procedure based on deep learning was performed to determine the genes that distinguish the normal tissue from the tumor tissue.

#### Materials and Methods

Gene expression data were taken from the GENT2 database (7). This database comprises the information on 68287 samples of patients’ tissues and cell lines, of which 58041 were tumor samples and 10246 were normal samples. We set the parameter of belonging to a tumor or healthy sample as the predictive target.

#### Results

Deep learning was performed according to the above described method with the ROC-AUC indicator of 0.83. 4912 genes were selected in the process of determining the significant features. The genes with the expression associated with the distinguishing a tumor tissue from the normal tissue were isolated from the previously found 36 ones with the help of the obtained list of genes. These 12 genes were: DKK4, GP6, LEF1, CUL1, KRT10, PIAS4, FSHR, MYLPF, EFNA3, ZFYVE9, GHSR, and MYL10.

### 3. NLP search for inhibitors

#### Introduction

To facilitate decision making on the choice of the revealed inhibitors, we designed a table with the help of python library Biopython that summarizes the results of a text search in PubMed PMC open source bases. Table 2 presents the number of articles published in response to various requests for each of 12 genes of interest. Such a table will help to draw a conclusion about the studying extent of the gene as a target.

**Table 2.** Results of NLP search for keywords associated with the studied genes and their inhibitors. Tags: green - low molecular weight inhibitor that triggers apoptosis, blue - protein, red - toxin

| Gene | Gene + 'target' | Gene + 'cancer' | Gene + 'lung cancer' | Gene + 'phase' | Gene + 'drug' | Gene + 'approval' | Gene + 'FDA' | Total mentions | Inhibitors |
| --- | --- | --- | --- | --- | --- | --- | --- | --- | --- |
| LEF1 | 9661 | 9138 | 4301 | 4962 | 4771 | 2129 | 665 | 35627 | imatinib,<br>wnt10b,<br>dlx3,<br>sb203580,<br>ex527,<br>dasatinib,<br>t0070907,<br>sb431542 |
| CUL1 | 4854 | 4245 | 1823 | 3629 | 2518 | 634 | 374 | 18077 | fbx4,<br>selumetinib,<br>fbxo7,<br>fbxo31,<br>fbxo21,<br>fbxo4,<br>fbxw7 |
| FSHR | 2143 | 2150 | 466 | 1657 | 1752 | 752 | 205 | 9125 | sunitinib, uk5099, clxbpa |
| GP6 | 1385 | 1977 | 460 | 1023 | 770 | 766 | 199 | 6580 | ono1714 |
| GHSR | 1308 | 926 | 365 | 828 | 1224 | 382 | 417 | 5450 | gefitinib, pd98059, pd90859, sb203580 |
| KRT10 | 934 | 998 | 388 | 700 | 604 | 320 | 75 | 4019 | r115866, sb431542 |
| PIAS4 | 810 | 792 | 319 | 505 | 427 | 127 | 53 | 3033 | nur77, pax8, foxm1b, trim32, zif268 |
| DKK4 | 652 | 661 | 339 | 312 | 368 | 139 | 48 | 2519 |  |
| EFNA3 | 542 | 541 | 328 | 272 | 318 | 141 | 28 | 2170 |  |
| MYLPF | 301 | 251 | 98 | 192 | 168 | 90 | 17 | 1117 |  |
| ZFYVE9 | 166 | 155 | 82 | 85 | 75 | 46 | 12 | 621 |  |
| MYL10 | 73 | 74 | 33 | 50 | 28 | 35 | 2 | 295 |  |

**Table 2.** Results of predicting the inhibitors for the target genes

| Gene | Number of inhibitors found |
| --- | --- |
| LEF1 | 639 |
| MYL10 | 524 |
| FSHR | 461 |
| EFNA3 | 385 |
| GHSR | 279 |
| CUL1 | 248 |
| GP6 | 190 |
| DKK4 | 134 |
| PIAS4 | 61 |

However, some references may not mean that there is a direct connection between the name of the gene and the drug used. In other words, they may be not related to it in terms of inhibition, but simply mentioned in a similar context.

#### Materials and Methods

Therefore, we developed a deep learning-based tool for the named entity recognition (NER) based on technology of natural language processing (NLP), for which we trained the NLP algorithm on the abstracts of articles labeled by hand. We identified the name of the gene or protein of interest and the name of its inhibitor. We used the BERT algorithm (8) as the basis. We applied the fine tuning procedure for this algorithm, which included training on the dataset of the labeled abstracts with BRAF gene and its inhibitors.

The created algorithm helped to achieve 98% accuracy of prediction.

#### Results

As a result, we added the right column to the table, which lists all inhibitors of a certain gene. These data allow a researcher to make a decision on the basis of the available number of inhibitors for each of the genes under consideration.

### 4. Drug-protein interactions

#### Introduction

Predicting effective interactions between a drug and a protein is a key to the search for candidate molecules. We have created a deep learning model with the initial data including both protein information and drug molecule information. The model is aimed at predicting the drug molecule effect on the protein.

The range of potential targets may shrink at this stage. This is due to the fact that not all of the genes selected at the previous stages can be qualitatively inhibited, or these gene products in the form of proteins may not bind effectively to the drug molecules.

#### Materials and Methods

We obtained the drug data from the open Drugbank database (9). This database comprises the data about drugs in combination with targets, including the protein that the drug is targeting, as well as structural representations of the molecules. We have selected only those small molecules for which there is a representation in the SMILES format. We needed two types of data to prepare the dataset: a target protein and a structural representation of the molecule.

As a result, the data array included positive examples with 19256 interactions for 5769 drugs and comprised 4104 unique proteins encoded by 3516 genes.

A challenge was to find negative examples for the training set. The researchers solve the problem in different ways: for instance, Wang and co-authors (10) reported that they randomly selected negative drug-target pairs with no interaction data. Researchers of another study also obtained negative examples extracting pairs with no interaction data, while randomly choosing the number of examples equal to the number of positive examples (11). Some authors predicted the absence of interaction by the algorithm (12). We analyzed the STITCH database (13), which contains scores of interactions between proteins and compounds.

To determine which score to regard as negative, we correlated interactions from the STITCH database with the Drugbank database, which included only pairs with a positive score. Thus, we expected we could understand which value to consider as a “positive speed”.

The left part of Figure 1 presents a boxplot for those pairs that are present in both databases (STITCH and Drugbank), therefore, they are considered positive. The right part presents a boxplot for all values from the STITCH database. Thus, it is evident that the range of positive rates does not intersect with the main range of data from the database of all interactions, and is an outlier for it.

**Figure 1.**
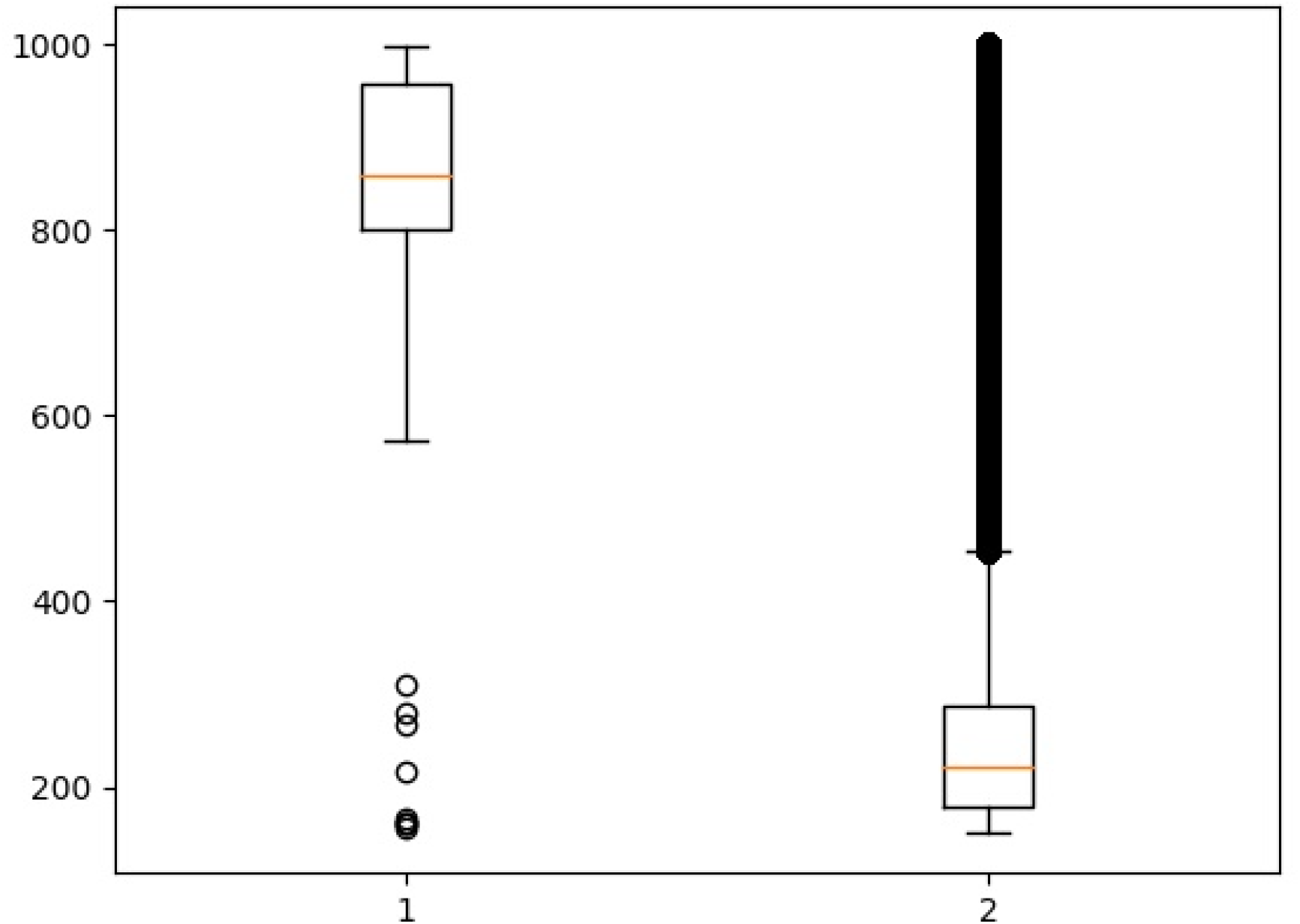
1 - The strength of the interaction between the compound and the protein, described in both databases (STITCH and Drugbank). 2 - boxplot for all values from the STITCH base.

**Figure 2.**
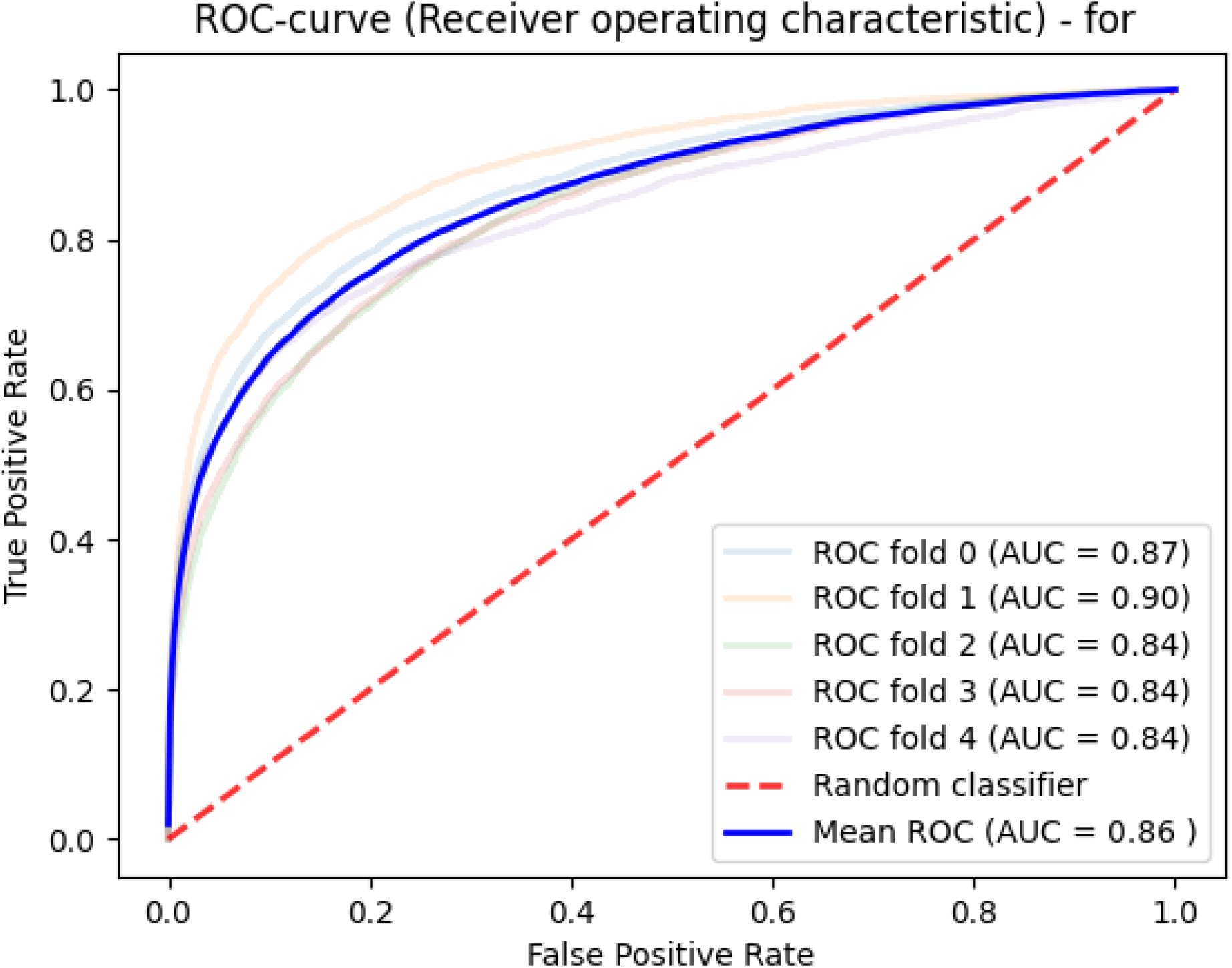
Model quality for predicting drug-protein interactions.

**Figure 3.**
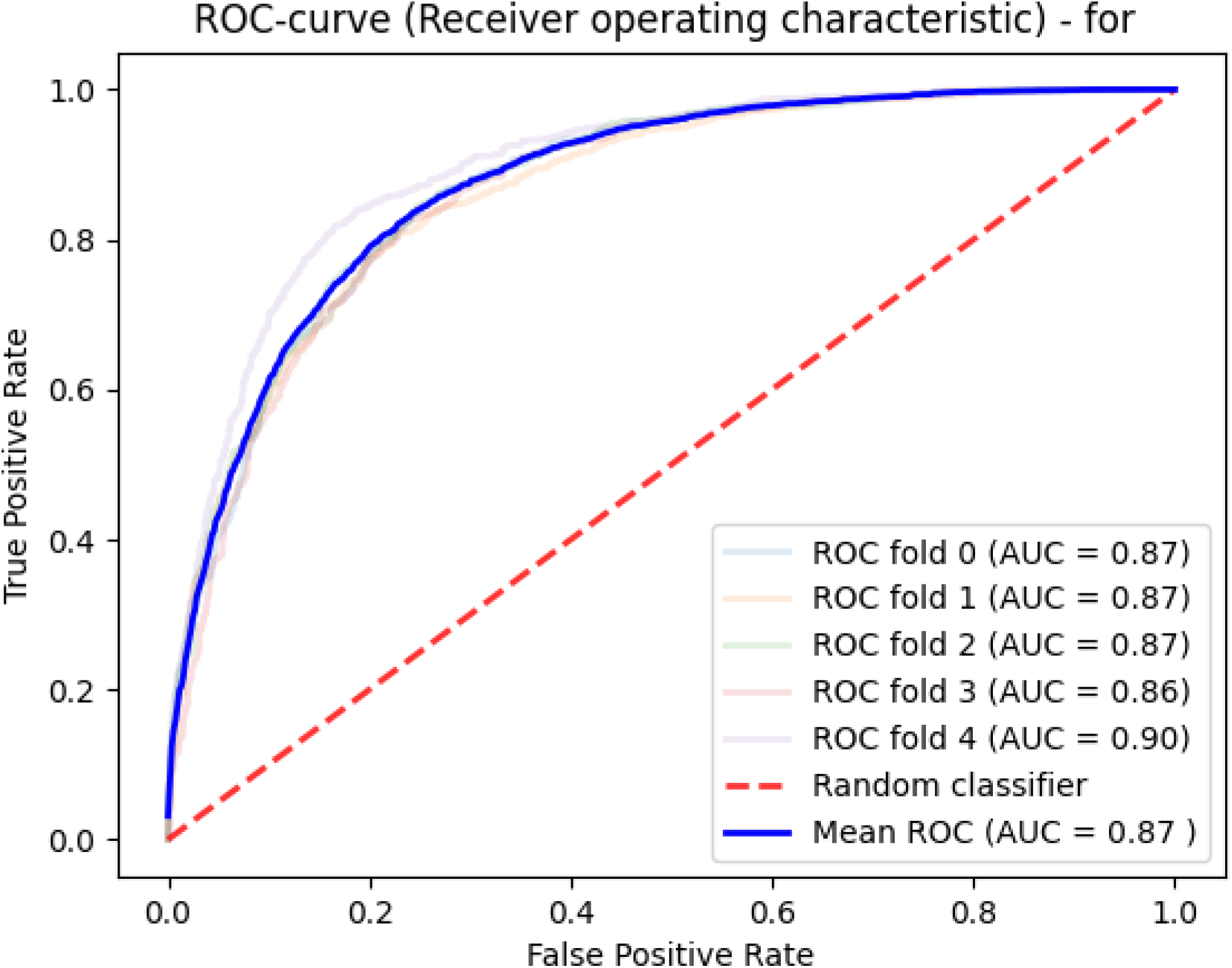
Quality of the model designed to assess the likelihood of reaching the compound IC50 data on lung cancer cell lines.

We used the lower quartile of interaction rates to form a sample of non-protein-binding drugs. We sorted the compounds of the obtained data according to the number of known interactions with proteins, and selected the 50,000 most common ones.

Amino acid sequences were obtained from the UniProt database (14). We presented them in vector form using the approach described elsewhere (15), where the authors vectorized all possible amino acid triplets (8 thousand) in the form of a 100-dimensional vector, and thus the vector representation of any protein consisting of these triplets was equal to their vector sum.

We also presented the compounds included in the dataset in a vector form, and in the form of 100-dimensional vectors, using the embedding approach of natural language processing technologies and implemented in the RDKit (16), mol2vec (17) and word2vec (18) libraries for python3 language.

As a result, a dataset was obtained from: 118379 pairs describing the compounds bound to proteins and 19250 pairs describing non-protein bound compounds (99129 precedents).

Deep learning was applied in a similar way as in the previous approach. ROC-analysis of learning quality allowed us to obtain an average area under the curve of 0.86.

To search for candidate molecules, we performed an experiment of predicting interactions for pairs of gene and compound of all possible ones.

We used all the molecules from the Pubchem library which had representations in the form of SMILES (23 million in total). They were presented in a vector form like that in the learning process.

Amino acid sequences of the encoding proteins were obtained from the UniProt database for 12 genes that we received earlier.

The prediction result was the DPI probability.

#### Results

We received 160,000 pairs with an interaction probability over 0.99, as well as 2921 pairs with an algorithm predicted probability of 1.0.

The following distribution by the inhibited genes was obtained for these 2921 potential inhibitors:

### 6. Prediction of the experiment on cell lines

#### Introduction

Emulation of a cell experiment is necessary in order to formulate the conditions for laboratory validation of the detected molecules, in particular, to determine cell lines on which preclinical trials will show a representative result. We performed a study for prediction of the chance for all compounds obtained at the previous step to reach IC50 concentration in cell lines.

#### Materials and Methods

The dataset was formed using the data of gene expression profiles of the cell lines from the Cancer Cell Line Encyclopedia (CCLE) database and compound sensitivity data for cell lines from Genomics of Drug Sensitivity in Cancer (GDSC) database (19, 20). We selected only lung cancer cell lines. A total of 11330 interactions were obtained for 122 drugs and 32 thousand genes in 106 cell lines.

The algorithm for determining the importance of the features selected 129 genes. The characteristics of the proteins encoded by the revealed genes are presented in Table 3.

**Table 3.**
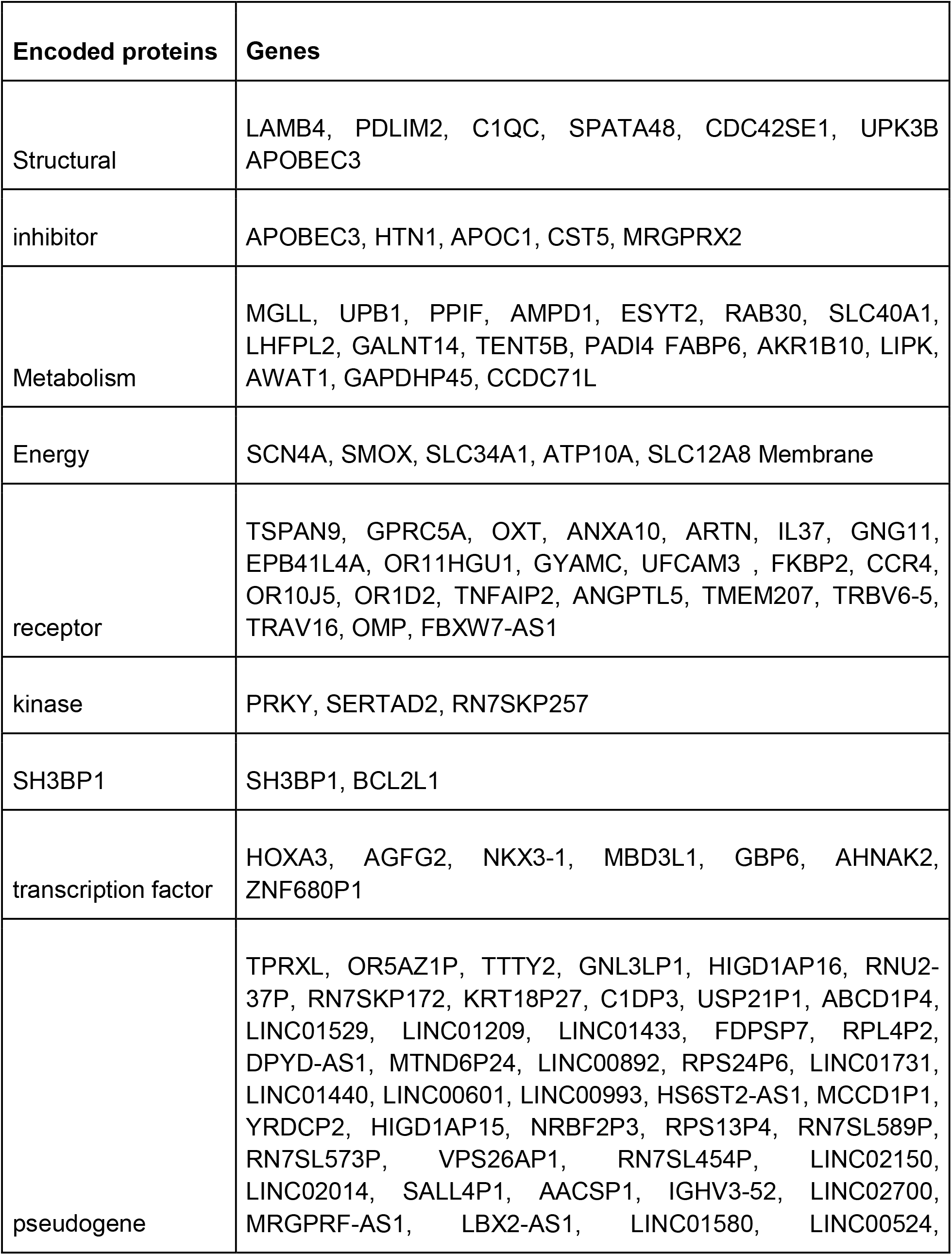
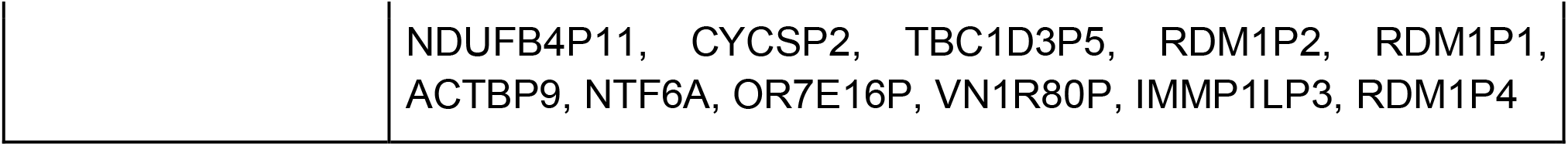
Characteristics of proteins encoded by the revealed genes.

Then, a study was performed for the prediction of these 106 lines interactions with potential inhibitors. We substituted each of the 2921 candidate inhibitors in turn, and predicted success of the IC50 test. After that, we selected all molecules with a probability of over 0.9 achieving IC50 from the results in the A549 and CALU1 cell lines. These cell lines were chosen due to the most frequent use of these lines in various studies of lung cancer.

The study resulted in the obtained data of the structure of 37 molecules with potential toxicity for lung cancer cells. We performed visualization in our own module, and found that the resulting molecules were large and consisted of many repeating structural elements of radicals. Therefore, we decided to isolate the active parts of the molecule for further analysis. As a result, another 15 molecules after their decomposition were added to the initial set of 37 molecules.

At the next step, we planned to test the selectivity of the obtained molecules in all 1018 cell lines. We designed a similar experiment to predict the IC50.

#### Results

We chose interactions with a probability of at least 0.9 from the data of the forecast. Molecules were selected that acted on the minimum number of lines with a probability higher than a given one, i.e., those with the highest specificity.

As a result, 5 small molecules were selected. The certain cell lines should be used for validation, such as ‘A549’, ‘NCIH23’, ‘NCIH460’, ‘NCIH1299’, ‘HCT116’, ‘AMO1’, ‘PC3’, ‘CAPAN1’.

## Conclusion

At the first stage of target discovery were selected about 36 genes. Interestingly, quite a few genes appeared to be associated with WNT signaling (DKK4, LEF1, WNT6, WNT8B), with BMP (SMAD9), and TGF (ACVR2A, GDF6, ZFYVE9). The study has found various components of the cytoskeleton and membrane proteins responsible for the transfer of various molecules. Potentially, each of these genes encodes proteins suitable for targeted lung cancer therapy.

Four of the genes found at the stage of tumor-normal filter, encode cytoskeletal proteins. Notably, the processes of tumor invasion and metastasis are often registered by the time of the diagnosis in patients with lung cancer (21). These features of the tumor are mediated by the developed cytoskeleton in the tumor cells. Thus, it is not surprising that there is a correlation between the expression of cytoskeletal proteins and a decrease in the overall survival of patients with lung cancer.

Natural language processing technologies used in the work have shown effectiveness for processing tens of thousands of articles. They can also be similarly used to compile own databases of scientific publications.

The pipeline of methods presented in this paper can serve as the basis for the technology of automated AI-driven drug discovery. The application of modern methods of machine learning, in particular, deep learning, as well as ways to present initial data for learning algorithms, is demonstrated. The performance of the methods, confirmed by cross-validation approaches on known results, was demonstrated using data from open sources. Ways to improve the methodology are the use of more data, including proprietary, as well as a more detailed representation of the original knowledge, in particular - 3-dimensional modeling of interacting molecules.

